# Internal timing-related dopaminergic dynamics can be explained by reward-prediction errors

**DOI:** 10.1101/2020.06.03.128272

**Authors:** Allison E. Hamilos, John A. Assad

## Abstract

Dopaminergic neurons (DANs) exhibit complex dynamics across a variety of behavioral contexts, often in ways that seem task-specific and even incompatible with results across different paradigms. Dopaminergic signaling during timing tasks has been a prime example. In behavioral timing, dopaminergic dynamics predict the initiation of self-timed movement via a seconds-long ramp up of activity prior to movement onset, similar to ramping seen in visuospatial reward approach and multi-step, goal-directed behaviors. By contrast, in perceptual timing, DANs exhibit more complex dynamics whose direction of modulation seems to be the *opposite* of that observed in behavioral timing. Mikhael et al. (2022) recently proposed a formal model in which dopaminergic dynamics encode reward expectation in the form of an “ongoing” reward-prediction error (RPE) that arises from resolving uncertainty of one’s position in the value landscape (i.e., one’s spatial-temporal distance to reward delivery/omission). Here, we show that application of this framework recapitulates and reconciles the seemingly contradictory dopaminergic dynamics observed in behavioral *vs* perceptual timing. These results suggest a common neural mechanism that broadly underlies timing behavior: trial-by-trial variation in the rate of the internal “pacemaker,” manifested in DAN signals that reflect stretching or compression of the derivative of the subjective value function relative to veridical time. In this view, faster pacemaking is associated with relatively high amplitude dopaminergic signaling, whereas slower pacemaking is associated with relatively low levels of dopaminergic signaling, consistent with findings from pharmacological and lesion studies.

## Introduction

Clues from human movement disorders and pharmacological studies have long suggested a connection between the neurotransmitter dopamine and the timing of movement initiation (Mikhael and Gershman, 2019; Soares et al., 2016; Dews and Morse, 1958; Schuster and Zimmerman, 1961; Meck, 1986; Rakitin et al., 1998; Lustig and Meck, 2005; Meck, 2006; Merchant et al., 2013; Malapani et al., 1998). We recently showed that dopaminergic signaling controls the moment-to-moment timing of movements in mice (Hamilos et al., 2021). We recorded dopaminergic signals with fiber photometry in mice executing a behavioral timing task in which animals had to self-time their movements to receive reward. Mice received juice for withholding movement for a proscribed interval (3.3 s) after a start-timing cue and then initiating movement (a first-lick) within a rewarded time window (3.3-7 s, Figure 1A).

**Figure 1:**
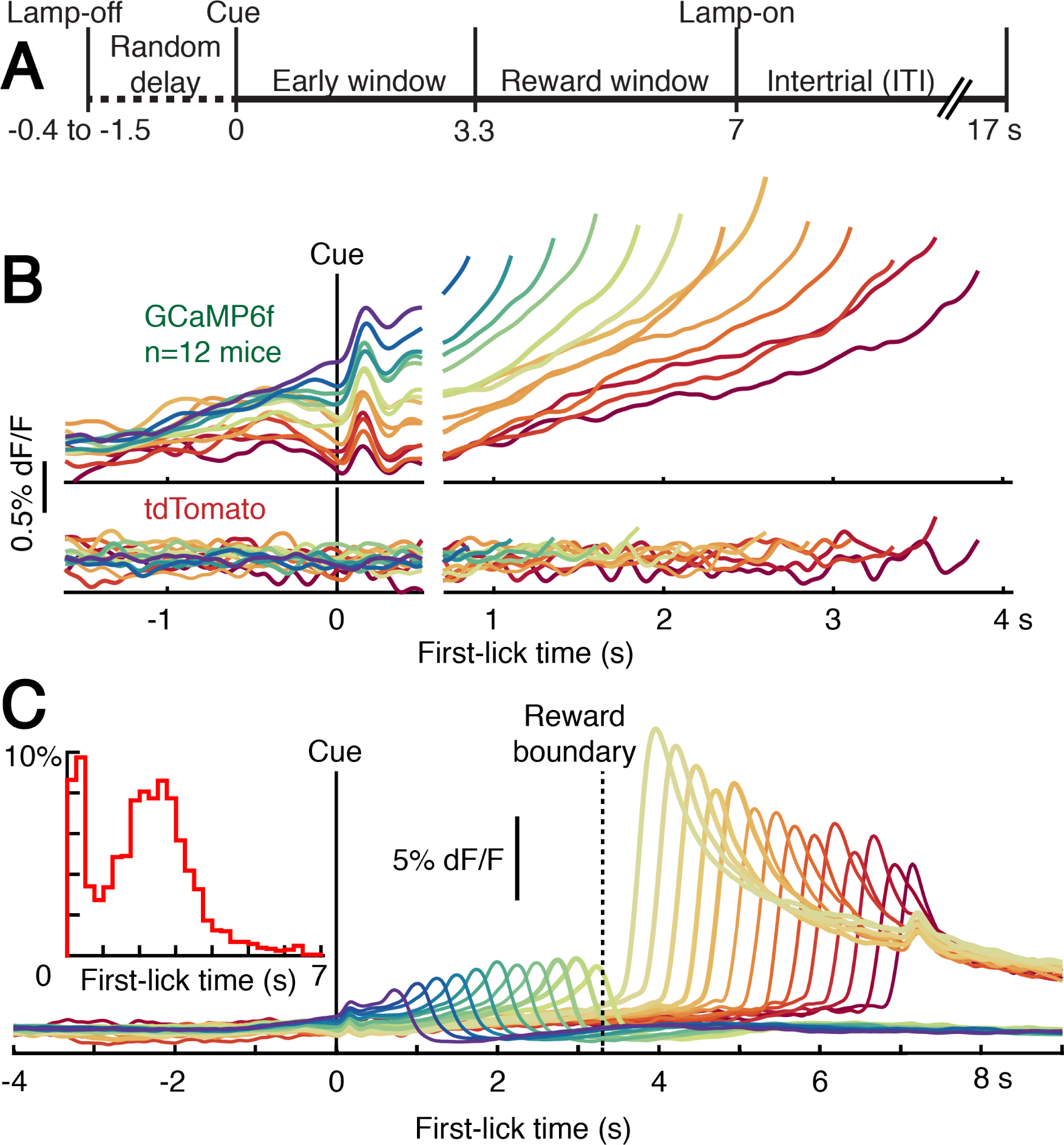
Nigrostriatal dopaminergic signaling during behavioral timing. Figure adapted from Hamilos et al. (2021) and used with permission. (A) Schematic of self-timed movement task. (B) Top: Average DAN GCaMP6f responses from 12 mice; Bottom: Responses of tdTomato, a non-activity-dependent fluorophore used to control for optical artifacts. The different colored traces correspond to averaged trial responses with different first-lick times (ranging from 1-4 s in 250 ms increments). Traces are plotted up to 150 ms before first-lick. Averaged traces are aligned relative to both start-timing cue onset (left of x-axis break) and first-lick (right of x-axis break); the break in the x-axis indicates the change in plot alignment. (C) Cue-aligned average DAN GCaMP6f signals at lower gain show post-movement RPE-like signals. Movement onset occurs just before the peak response for each curve. Mice were rewarded for first-licks made later than 3.3 s, but were not rewarded for earlier first-licks. Inset: distribution of first-lick times, n=12 mice.

We observed two aspects of dopaminergic signaling that predicted movement timing: 1) pre-trial baseline signaling of nigrostriatal dopamine neurons (DANs), and, 2) slow “ramping” signals that built up over the course of seconds between the start-timing cue and the self-timed movement (Figure 1B). Although self-timed movements occurred with variable timing relative to the start-timing cue (Figure 1C, inset), DAN signaling rose to about the same level at the moment of movement onset, reminiscent of a ramp-to-threshold process. DAN signals were not explained by ongoing nuisance movements nor optical artifacts and were best modeled with timing-dependent predictors, including a baseline offset term whose amplitude was proportional to the mouse’s timing on the upcoming trial, as well as a stretch feature that encoded percentages of elapsed time between the cue and self-timed movement (Hamilos et al., 2021). DAN ramping activity predicted first-lick time on single trials, independently of trial history, and optogenetic manipulation of DANs bidirectionally shifted movement timing, with activation early-shifting movements versus inhibition late-shifting movements. Together, these results indicate that dopaminergic signaling during self-timing controls the moment of movement onset.

### A temporal-difference learning model of dynamic dopaminergic signaling

We were interested in understanding the origin of the dynamic dopaminergic signals we observed in our self-timed movement task and how they fit into the context of prior work on the dopamine system. A framework that has explained many disparate experimental results from the dopaminergic system is temporal difference (TD) learning with reward-prediction errors (RPE; Mikhael et al. (2022); Schultz et al. (1997)). In this framework, DAN activity is thought to reflect the moment-to-moment difference in the animal’s expectation versus its perception of the value of its current state, where value is defined as the temporally-discounted expectation of total future reward. In classical trace-conditioning paradigms, DANs fire in transient bursts to unexpected rewards and reward-predicting cues, whereas they pause their firing when expected reward is omitted. Indeed, we observed RPE-like signals in the cue-related transient, dips in activity after unrewarded first-licks, and surges in activity following rewarded first-licks (Figure 1C). Persistence of RPE-like signals in well-trained animals has been suggested to arise from the inherent imprecision in neural timing (Rakitin et al., 1998), which may reflect the animal’s moment-to-moment uncertainty of its current state—*i.e*., its position in time–and, by extension to our task, uncertainty about its accuracy for a given self-timed movement (Mikhael and Gershman, 2019). Indeed, positive-going RPE-like signals were strongest for first-licks closest to the reward-boundary (3.3 s), presumably when the mouse’s “confidence” of reward was lowest, consistent with the greatest RPE occurring when the mice were least certain of success (Figure 1C).

Whereas RPE-frameworks have explained *transient* bursts and pauses in DAN activity during trace conditioning and other types of learning experiments, DAN activity can also change more slowly (Mikhael and Gershman, 2019; Mikhael et al., 2022). For example, “ramping” signals build up over seconds during goal-directed navigation (Howe et al., 2013), bandit tasks in which animals must complete multiple goals to receive reward (Mohebi et al., 2019; Hamid et al., 2016), and tasks with visual cues of proximity to reward (Kim et al., 2019). It has been suggested that DANs could signal different information via slow changes in activity (*e.g*., motivation, ongoing value, vigor) compared to fast-timescale activity (*e.g*., post-hoc RPE signals for learning), and a number of proposals have suggested that DANs multiplex different kinds of information over different timescales and contexts (Mohebi et al., 2019; Engelhard et al., 2019).

However, recent models have proposed RPE-based explanations that may be able to reconcile these seemingly disparate dopamine signals (Mikhael and Gershman, 2019; Mikhael et al., 2022; Kim et al., 2019). While these models do not preclude the possibility that DANs could also encode other types of information (*e.g*., value, vigor, *etc*.), they are attractive for their parsimonious explanation of how fast timescale phenomena and slowly-evolving ramps could arise from the same underlying RPE-based calculation. In short, these models employ principles from TD learning to show how certain shapes of the value function (*i.e*., the assignment of values to the series of behavioral states comprising a task) can give rise to a *continuously changing* RPE, even in well-trained animals (Mikhael and Gershman, 2019; Mikhael et al., 2022; Kim et al., 2019; Starkweather et al., 2017).

We were interested in whether an RPE-based framework could explain the results found in our self-timed movement task as well as results from other timing tasks (Soares et al., 2016). To approach this question, we applied a key feature of TD learning algorithms to determine what an RPE-like signal would look like in different kinds of timing tasks. Specifically, we took advantage of the fact that *RPE is proportional to the derivative of the subjective value function under conditions of state uncertainty* (Mikhael and Gershman, 2019; Mikhael et al., 2022), as is the case during timing tasks in which the animal must rely on its own internal representation of time to guide behavior (Mikhael et al., 2022).

Thus, if the value landscape for a given behavioral task is known, and if DAN activity encodes RPE, the RPE-based framework makes predictions about the expected shape of dynamic DAN activity during the task. In a recent study, similar applications of this principle predicted the ramping DAN signals that were observed in virtual reality (VR) tasks in which animals were moved passively through VR spaces, as well as when the animals passively viewed abstract, dynamic visual cues indicating proximity to reward (Kim et al., 2019; Mikhael et al., 2022), suggesting that the ramping in our task could be explained from similar principles.

## Results

### Simulation of Temporal Difference Learning in behavioral timing

In a simple TD learning model of self-timed movement, time may be modeled as a continuous set of states through which a Markov agent must traverse to receive reward (Sutton and Barto, 1998) (Figure 2A). At each state transition (timestep), the agent must decide whether to move (lick) or to wait based on the probability of transitioning to a reward or failure state. If the agent is an optimal timer, its subjective approximation of its current state, *τ*, accurately tracks veridical time, *t*, and it will thus withhold movement until the first moment at which reward will be available in response to licking (3.3 s in our self-timed movement task, Figure 1A).

**Figure 2:**
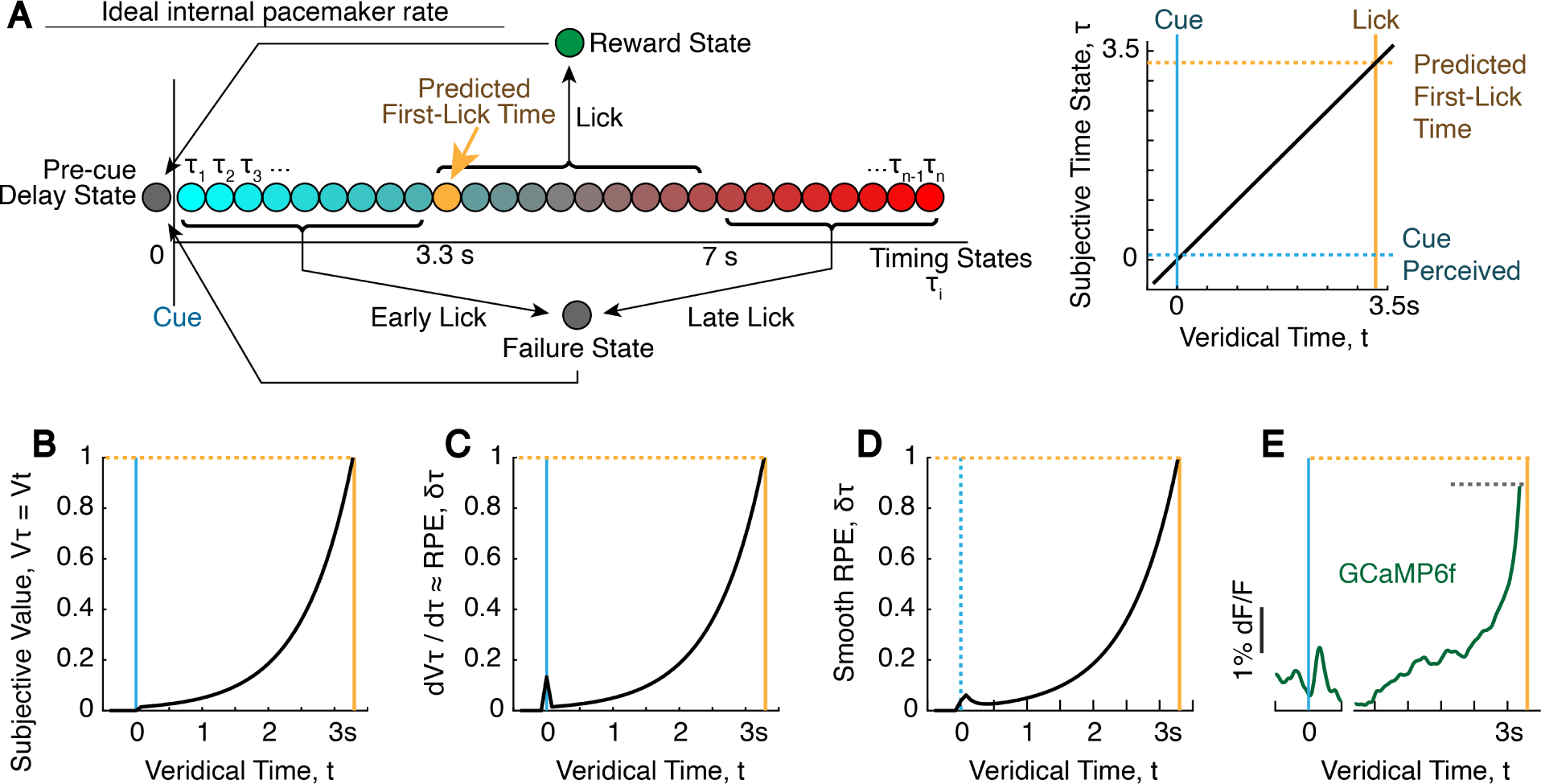
Simulated Value and RPE functions for an optimal timer during behavioral timing. (A) State space and probability of state transition for an optimal timer. Gold-shaded state is the first state from which reward is available, and thus is when the first-lick is predicted to occur. (B) Simulated learned estimated value function 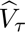 for an optimal timer, where 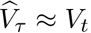, assuming conditions of self-resolved state uncertainty (Mikhael et al., 2022). The agent is expected to first-lick at the peak of the trajectory (gold). The learning model produces an exponential value landscape. However, any sufficiently convex value function could produce ramping RPE functions (Mikhael and Gershman, 2019; Mikhael et al., 2022; Gershman, 2014). (C) Simulated learned RPE function for an optimal timer. The learned 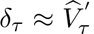 the derivative of the subjective value function. (D) Predicted DAN GCaMP6f signals for an optimal timer if DAN signals reflect RPE. The RPE function was smoothed with a causal filter to approximate GCaMP6f off-kinetics. (E) Average SNc DAN GCaMP6f dynamics (n=1 mouse, 5 sessions) shown for comparison. First-lick time: 3.33-5.1s (chosen to match timespan in Figure 3); trace plotted up to 150ms before first-lick to exclude potential movement artifacts (grey dashed line). Break in x-axis as in Figure 1B.

The value landscape for this model of the self-timed movement task can be understood intuitively: When the cue event occurs, a well-trained agent can expect an increased possibility of reward in the next few seconds; thus, at this moment, value increases. However, reward never occurs within the first 3.3 s of the standard timing task we implemented; thus, value at the cue is necessarily lower than value at 3.3 s. In fact, value will continually increase as time approaches 3.3 s. Thus, as long as the agent withholds movement, the value landscape, *V_t_*, during the first few seconds is a monotonically increasing, convex function (Gershman, 2014) (Figure 2B).

However, even an optimal timer does not have access to the true state identity, *t*. Consequently, it can never be certain of its subjective approximation of its state, *τ*. If this uncertainty is not resolved (such as in Pavlovian tasks), the learned value function will be distorted, and RPE signals are not expected to ramp (Mikhael et al., 2022). This would be the case in a typical Pavlovian task, in which reward is delivered after a fixed delay relative to a cue, regardless of the agent’s behavior.

But behavioral timing creates a different scenario: unlike Pavlovian conditioning, there is an onus on the agent in behavioral timing to *act* to obtain reward. This requires that the agent must *somehow* resolve its state uncertainty to decide when to move.

Intriguingly, it was recently shown that when animals have visual cues indicating their proximity to reward in Pavlovian conditioning, dopaminergic ramps *do* occur (Kim et al., 2019). Mikhael et al. (2022) proposed a model to explain this phenomenon, in which the visual cues allow the animal to resolve its state uncertainty. This resolution of uncertainty would provide an update to the animal’s distorted estimate of the value function at each moment in time. As a result of this update, there would be non-zero RPE at each moment of time, resulting in a dynamic RPE function. In their model, this dynamic RPE function is expected to be approximately the derivative of the subjective value function (Mikhael et al., 2022; Kim et al., 2019), 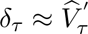, where *δ _T_* s RPE at subjective time *τ*, and 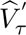 is the time-derivative of the subjective value function.

Because agents must somehow resolve their state uncertainty to act in behavioral timing, we suspected a similar phenomenon occurs in these tasks. However, because information is not available from the environment, agents would instead have to *self-resolve* their own state uncertainty by relying on their estimate of their time state. If agents act under this assumption (*τ ≈ t*), then state uncertainty could be self-resolved similarly to resolution with visual cues. We thus applied the value function learning model proposed by Mikhael et al. (2022) to simulate the RPE function in behavioral timing assuming self-resolution of timing state uncertainty.

The resulting shape of the RPE function, *δ_τ_* is simple to intuit: a transient increase at the cue followed by a slowly-evolving ramp (Figure 2C). If the agent is an optimal timer, the subjective estimate of the time state is consistent with veridical time (*τ* = *t*), and thus the learned value function exactly matches the true value function (Figure 2B). Thus, the agent is expected to move right at 3.3 s. If neural signals represent value or RPE functions, measuring them with a calcium indicator such as GCaMP6f would distort them due to the binding kinetics of the indicator. We thus simulated the expected GCaMP6f readout of the RPE and value functions by applying a causal filter to approximate indicator kinetics (Figure 2D).

The modeled RPE function mirrors the shape of the dynamics observed in DAN signals: a cuerelated transient followed by a slow ramp up to the time of first-lick (Figure 2D-E). However, unlike the optimal timer in this model, mice, like humans, exhibit suboptimal timing behavior with variability proportional to the duration of the timed interval (Rakitin et al., 1998). It has been proposed that this variability in timing results from imprecision in an internal clock, referred to classically as the internal “pacemaker” (Gibbon et al., 1997). When the pacemaker is fast, selftimed movements occur relatively early, whereas when the pacemaker is slow, later movements occur. These changes in the pacemaker rate would correspond to the mouse traversing the set of subjective states, *τ*, at different rates than the passage of veridical time, *t* (Figure 3A), resulting in relative *compression* or *stretching*, respectively, in the subjective value function, 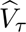 (Figure 3B), with corresponding compression/stretching of the RPE function (Figure 3C-D).

**Figure 3:**
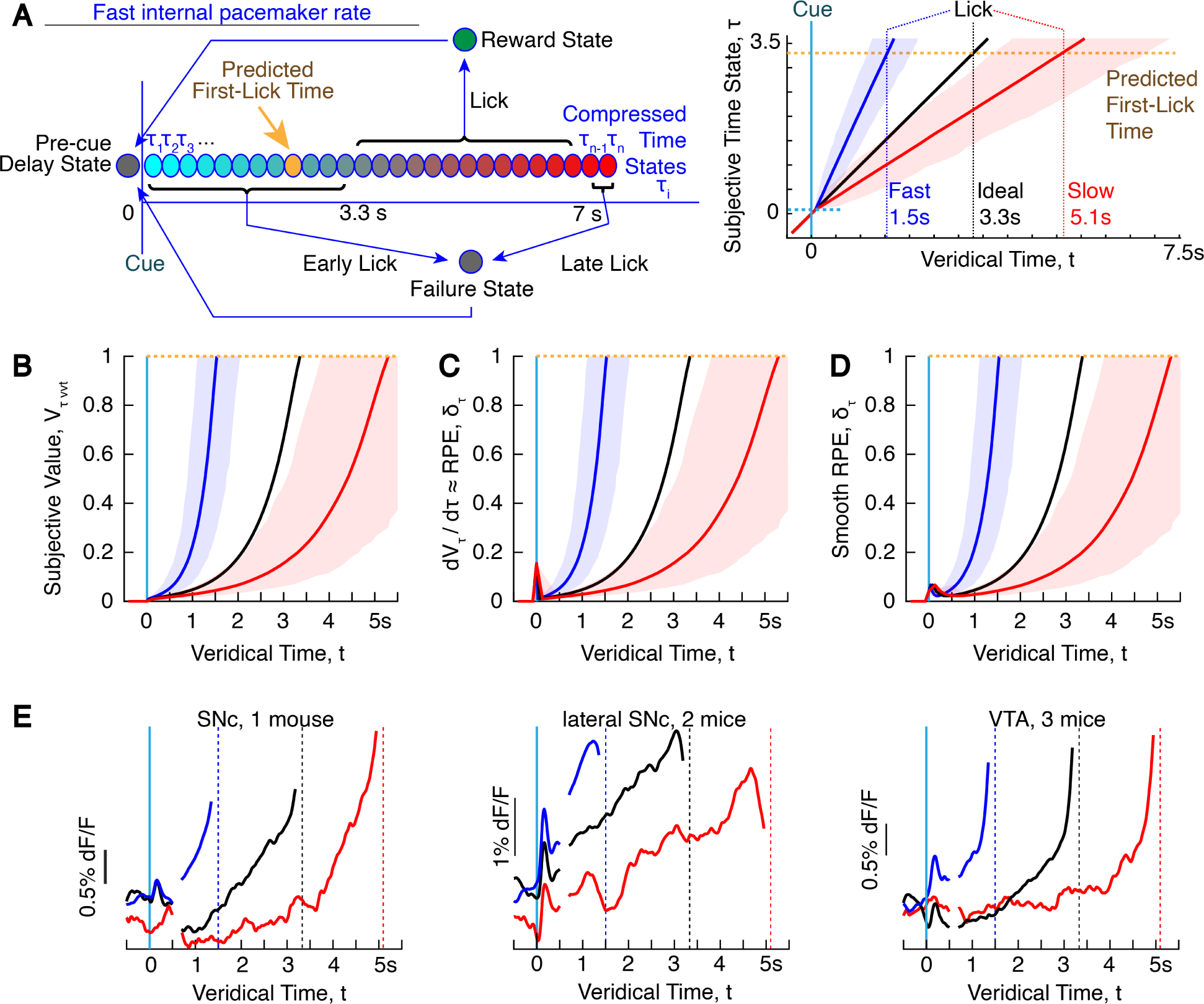
Simulated compressed/stretched Value and RPE functions for a sub-optimal timer. **(A)** Left: State space of self-timed movement task for a suboptimal timer with a fast pacemaker. The fast pacemaker “compresses” state space Mikhael and Gershman (2019); Sutton and Barto (1998), resulting in traversal of the timing states faster than veridical time. The mouse can only make a decision based on which state it believes itself in; thus first-lick is expected to occur early (gold-shaded state). Right: 1000x simulated state space trajectories for slow and fast internal clock. Shading: 95% confidence interval. **(B)** Simulated learned value functions. Fast pacemaking compresses the subjective value function (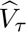, blue). Slow pacemaking stretches the value function (red). The animal is expected to lick at the peak of the trajectory. **(C)** Simulated learned RPE function and **(D)** its smoothed estimate, (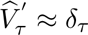). **(E)** Different shapes of dopamine ramp compression/stretching, plotted for the movement time ranges corresponding to panel A. *Left* : SNc GCaMP6f, 1 mouse, 4 sessions. (All 12 mice shown in Figure 1B.) *Middle*: SNc GCaMP6f signals from the 2 most laterally implanted mice. The linear ramping shape was consistent between these 2 mice and similar to signals recorded from DAN terminals in dorsolateral striatum (n=7 mice). *Right* : VTA DAN GCaMP6f signals (3 mice, 16 sessions). In general, there appeared to be a gradient of ramping shapes from more medial to lateral dopaminergic areas. Lateral SNc had more linear and down-turning ramps. VTA had more convex, up-turning ramps. Intermediate areas of SNc showed a mixture of both features, with less convexity and less pronounced up-turns.

Strikingly, as this simple RPE-based model predicts, DAN signals observed during our self-timed movement task show different ramping dynamics depending on when the animal actually moved (Figures 1B and 3E), consistent with compression/stretching of the subjective value and RPE functions. When the animal moved relatively early (perhaps corresponding to a fast pacemaker), DAN ramping unfolded with a steeper slope, as if the ramping period were *compressed*. Conversely, when the animal moved late (perhaps corresponding to a slow pacemaker), DAN ramping unfolded with a shallower slope, as if the ramping interval were *stretched*. The idea of compression/stretching of DAN ramps was supported by generalized linear encoding models (Figure 5C of Hamilos et al. (2021)), for which we needed to add a timing-dependent “stretch factor” to best capture the variance in GCaMP6f signals during the timed interval. Together, these observations could be explained by DANs encoding an RPE-like signal related to the animal’s “belief” of its position in objective time, *τ*, as derived from its position along the subjective value trajectory during the timing interval of the task.

In fact, a recent model described how a timing mechanism instantiated by the nigrostriatal system could lead to (the well-known) variability in self-timed intervals by stretching or compressing of subjective value trajectories (Mikhael and Gershman, 2019). The model posits that dopamine modulates the pacemaker rate (consistent with pharmacological and lesion studies), with increased dopamine availability (or efficacy) speeding the pacemaker, and decreased dopamine slowing the pacemaker (Dews and Morse, 1958; Schuster and Zimmerman, 1961; Meck, 1986; Lustig and Meck, 2005; Meck, 2006; Merchant et al., 2013). In turn, the pacemaker controls the encoding of subjective time, and thus the steepness of the value function with respect to objective, veridical time. It follows that variation in dopamine availability would compress or stretch the value landscape to varying degrees from trial-to-trial. This model is consistent with our findings of variable ramping slope in DAN signals from trial-to-trial. It is also consistent with neural recordings from striatal spiny projection neurons and parietal cortical neurons during similar self-timed movement tasks, for which temporal sequences of striatal and cortical firing during timing were compressed for early movements and stretched for late movements (Mello et al., 2015; Maimon and Assad, 2006).

While the RPE-based view of DAN activity captures the dynamic DAN signals we observed, our simple RPE model alone does not capture the *baseline offsets* in DAN signals that were predictive of movement timing even after controlling for previous trial outcome and ongoing nuisance movements (Hamilos et al., 2021). More complex RPE-based explanations for these *tonic* offsets in DAN signals could be imagined with further assumptions (*e.g*., states like the pre-cue delay could also contain timing states that create offsets before the trial begins, *etc*.), but a parsimonious explanation for how and why these offsets emerge requires further investigation. Mohebi et al. (2019) recently showed baseline differences in the amount of dopamine in the nucleus accumbens core that were correlated with the recent history of reward rate: higher recent reward rates were related to higher tonic dopamine (Mohebi et al., 2019). However, in our task, animals tended to move later toward the end of sessions, resulting in periods of relatively high reward rate when the average tonic baseline signal was *lower* (baseline preceding rewarded trials—by definition, later movements—was systematically lower in our task, Figure 1B), suggesting a more complex relationship between tonic DAN activity and reward rate in our task. While the origin of offsets in DAN signals remains unclear, these offsets were nonetheless inversely related to the first-lick time, and thus directly related to the (inferred) pacemaker rate, consistent with pharmacological and lesion studies positing a positive correlation between dopamine availability and pacemaker rate (Mikhael and Gershman, 2019; Gershman, 2014; Soares et al., 2016; Dews and Morse, 1958; Lustig and Meck, 2005; Meck, 2006; Merchant et al., 2013).

### RPE-based predictions for DAN signalling during perceptual timing

Whereas DAN signals during behavioral timing observed in our self-timed movement task were consistent with classic observations of the influence of dopamine on the speed of the pacemaker, a recent study found more complex DAN dynamics during *perceptual* timing. Soares et al. (2016) recorded SNc DAN GCaMP6f signals with fiber photometry as mice executed a classic temporal bisection perceptual task (Soares et al., 2016) (Figure 4A). Trials began when mice entered a nosepoke port and received an auditory start-timing cue. Mice had to remain in the port throughout a variable timing interval, which was terminated with a stop-timing auditory cue. Mice then reported whether the interval was shorter or longer than a criterion time (1.5 s) by choosing a left or right nose-poke port corresponding to a “long” or “short” judgment. Mice were trained to categorize intervals spanning 0.6-2.4 s. As expected, trials with more extreme intervals were easier for the mice, whereas trials with intervals closer to the 1.5 s criterion time elicited chance performance (Figure 4B).

**Figure 4:**
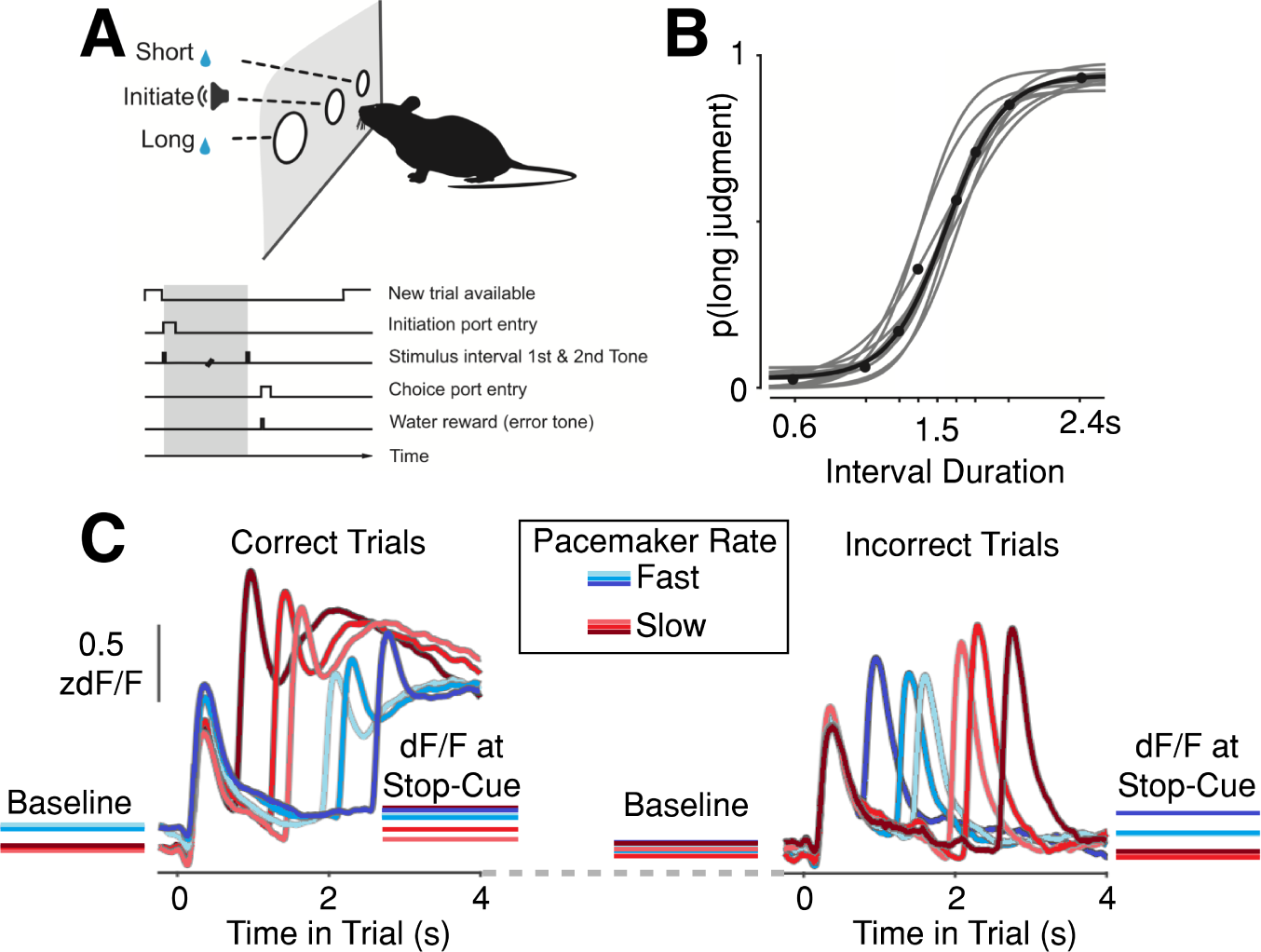
Complex dopaminergic dynamics during perceptual timing. Figures adapted with permission of authors from Soares, S., Atallah, B. and Paton, J. Midbrain DA neurons encode perception of time. *Science* **354**,1273–1277 (2016). Reprinted with permission from AAAS. (A) Task schematic. (B) Psychometric curve for timing intervals of different duration. Criterion time: 1.5 s. (C) Start-timing cue-aligned average SNc DAN GCaMP6f signals. Second peak occurs just after the stop-timing cue (intervals: 0.6, 1.05, 1.26, 1.74, 1.95, 2.4 s). Figure recolored to indicate average inferred pacemaker rate. Red: slow; blue: fast. Note: colors intended to indicate *category* of clock speed, ***<u>not</u>*** relative pacemaker speed within category. Left: Correct trials. Right: Incorrect trials.

DANs exhibited complex dynamics during the bisection task, starting with a sharp transient after the start-timing cue and ending with a second transient after the stop-timing cue (Figure 4C). Between the start-timing and stop-timing cues, DAN signals exhibited a U-shape with increasing time, which was visible for trials with longer intervals but was truncated prematurely for the shorter intervals. The authors focused their analyses on the transient occurring *after the stop*-*timing* cue. Short judgments (suggesting a slow pacemaker) were accompanied by relatively high-amplitude transients after the stop-cue, whereas long judgments (suggesting a fast pacemaker) showed relatively low-amplitude transients. These results seemed to suggest that relatively *high* DAN activity reflected a *slow* pacemaker, the opposite of what is expected based on the bulk of pharmacological and lesion studies (Mikhael and Gershman, 2019), as well as the trend we observed during our self-timed movement task.

This surprising finding could be a unique feature of the bisection task. Unlike self-timed movements, in which animals directly report elapsed time with a movement, the temporal bisection task requires an additional computational step, in which the timed interval must be categorized as “long” or “short.” However, prior pharmacological studies employing the bisection task found results consistent with the classic view that higher dopamine availability is associated with a faster pacemaker (Mikhael and Gershman, 2019; Morgan et al., 1993)—opposite the interpretation of Soares et al. (2016), but consistent with the findings of our self-timed movement task.

The discrepancy between our results and those found by Soares et al. (2016) could perhaps be traced to differences in the way DAN signals were analyzed. We focused our attention on DAN signals unfolding *during timing* in our self-timed movement task, whereas these signals were not explored by Soares et al. (2016). We thus asked two questions: 1. What correlations exist between DAN signals and pacemaker rate in the bisection task *before* the timing interval? And, 2. What correlations exist *during* the timing interval itself?

Before addressing these questions, we note that the relationship between pacemaker and bisection judgment is not as straightforward as in self-timed movement, and thus we recolored Figure 4C to clarify this, employing the following intuition: For a trial to be correct in the bisection task, on average, the pacemaker must be either accurate or “conservatively inaccurate.” In other words, a correct “short” judgment requires either accurate timing or a *slow* pacemaker (Figure 4C, red curves). Conversely, a correct “long” judgment requires either accurate timing or a *fast* pacemaker (Figure 4C, blue curves).

When we considered DAN signals *before* the timing interval for correct trials in the Soares et al. (2016) study (Figure 4C, left), we noticed what appears to be two strata of signal levels. Trials with “long” judgments (fast pacemaker on average) had relatively high baseline signals, whereas trials with “short” judgments (slow pacemaker on average) had lower baseline signals, consistent with the relationship between baseline offsets and pacemaker rate that we observed in our self-timed movement task. As in our task, these baseline offsets remained present during the timing interval, resulting in the same stratification of dF/F signals immediately prior to the stop-timing cue (except for the very shortest interval, 0.6 s, which overlaps decaying GCaMP6f signals related to the start-timing cue, likely causing an artefactual inflation of the signal just prior to the stop-cue due to the off-kinetics of the calcium indicator or kinetics of calcium clearance more generally). Thus, it generally appears that DAN activity was *higher* on trials with fast pacemaker rates, both during and before the interval in which the animal was actually timing. Intriguingly, *incorrect* trials (to the right in Figure 4C) showed a relative convergence of the baseline signals preceding the start-cue, but then signals diverged during the timing interval, resulting in relatively *high* signals just before the stop-cue for incorrect “long” choices (*i.e*., a fast pacemaker, blue), but relatively low signals just before the stop-cue for incorrect “short” choices (*i.e*., a slow pacemaker, red). This is consistent with the patterns observed on correct trials. Interpreted thusly, the Soares et al. (2016) result is consistent both with our behavioral timing results and with classic pharmacological studies relating higher/lower dopamine availability to faster/slower pacemaker rates, respectively. Soares et al. (2016) presented their subsequent analyses with these baseline differences normalized-out in some way (Figure 3 of Soares et al. (2016)). It is possible that this “zeroing out” of the baseline offset may have hindered efforts to detect consistent effects during the timing interval due to reordering of the traces.

Because baseline offsets in the bisection task appear similar to those in our self-timed movement task, we asked whether dynamic DAN signals in the bisection task could similarly be explained by the task’s RPE landscape. In their investigation of the stop-timing cue-related transient, Soares et al. (2016). showed that its amplitude is well-explained by a combination of temporal surprise and behavioral performance, and we applied these parameters to derive a value landscape consistent with their perceptual timing task.

The inferred value landscape of the bisection task for an optimal agent was built from a few assumptions (Figure 5A):

1. As in our self-timed movement task, value increases immediately at the start-cue and continues to rise toward the time of expected potential reward delivery.
2. Because the longest interval is 2.4 s, the time until potential reward is known to be no more than *∼*3 s (including the time to report judgment). However, due to temporal uncertainty and the fact that a false start (leaving the port before the stop-timing cue) results in an error and loss of reward, there is a second jump in the value function at the time of the stop-cue when the feedback of the tone reorients the value function and indicates the opportunity to collect reward within a few hundred milliseconds.
3. Because value is temporally discounted at the start-cue by the possibility of the longest-possible interval, any stop-cue occurring before 2.4 s results in a sudden “teleportation” through the value landscape to the final limb of the task that occurs just before the judgment and ascertainment of trial outcome, similar to the jump in the value function in a recently-reported, virtual reality, spatial teleportation task (Kim et al., 2019). Thus, assuming the value function trends upwards steadily, the amplitude of RPE-related transients following the stop-cue would *decrease* as the interval duration increases, because the sudden jump in the value function becomes progressively smaller.
4. To capture aspects related to behavioral performance, we additionally included contours in the value function during the timing interval to reflect the probability of a correct choice for intervals of different lengths. Specifically, a relative minimum in the value function occurs near 1.5 s, when predicted performance is worst. However, a stop-timing tone near the criterion time also results in a smaller jump in the value function because the probability of a correct decision is also lower. Thus, the increase in value at the moment of decision was adjusted by the probability of a correct choice.
5. As in the simple RPE-model of our self-timed movement task, we modeled changes in pacemaker rate as compression/stretching of the subjective value landscape with respect to veridical time.
6. The agent traverses timing states during the timing interval, similar to the timing states in the self-timed movement task, but unlike our task, the bisection task does not require the agent to decide when to move. We assume the need to make a timed movement imposes a need for the agent to be relatively certain of its subjective timing state, *τ*, to make a decision, even though it is uncertain of its true state, *t*. The bisection task, on the other hand, is more similar to classical conditioning tasks in which the timing interval is not in the agent’s control, and thus subjective state uncertainty increases with the distance from the last state-informative cue (Mikhael and Gershman, 2019). Thus, we took into account temporal blurring of the subjective state function, which would tend to reduce the convexity of the subjective value function and reduce the amplitude of ramping during the timing interval (Mikhael and Gershman, 2019). However, adding temporal blurring does not substantially change the fit-shape in our simplified model, and versions with or without blurring can reproduce the shape of the dynamic DAN signals.

**Figure 5:**
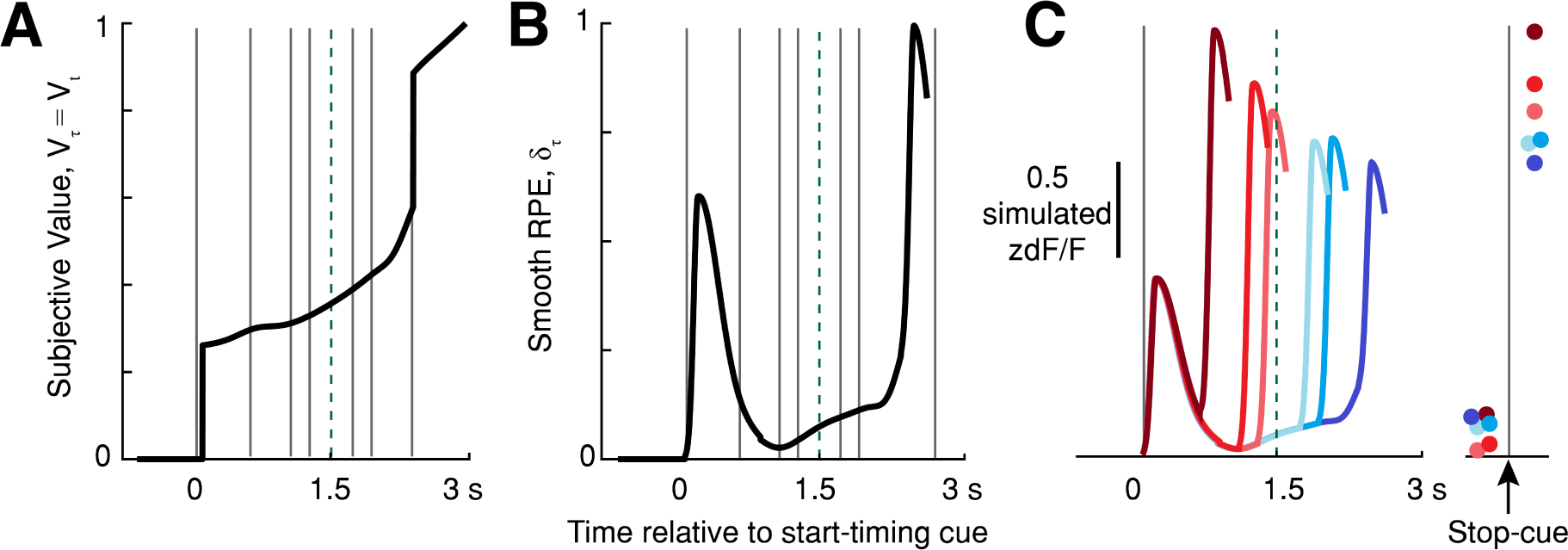
Subjective Value and RPE Landscapes for perceptual timing (bisection task). (A) Estimated value function 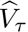, where 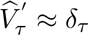 for an optimal timer on a 2.4 s trial. Grey lines: test interval times. Green dashed line: criterion time (1.5 s). Value increases approaching the time when reward is available, increasing abruptly at the start- and stop-timing cues (0 and 2.4 s). (B) Smoothed RPE function for an optimal timer, estimated as 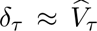, the derivative of the subjective value function. The RPE function was smoothed with an asymmetrical Gaussian kernel spanning *ca*. 28% of the interval to approximate GCaMP6f off-dynamics. (C) Predicted DAN GCaMP6f signals for an optimal timer for the six test interval times. Traces truncated before reward collection for clarity. Colors indicate conservative pacemaker speed for a correct judgment (red: slow, blue: fast). Right: relative simulated dF/F amplitude just before the stop-timing cue and subsequent peak response.

Together, we arrived at a model of the RPE landscape for each of the six tested interval durations (Figure 5B-C). Importantly, this simple RPE-based model accurately captures the relative categorical amplitudes of the stop-timing cue-related transients, as follows: If the instantaneous DAN activity at the time of the stop-timing cue is relatively high, this would indicate that the animal is further along in the subjective value trajectory, resulting in 1) a *long* judgment, and 2) a relatively *smaller* RPE transient, because the underlying subjective value was higher at that moment. Conversely, if instantaneous DAN activity is relatively low at the stop-timing cue, this would indicate that the animal is not very far along the subjective value trajectory, leading to 1) a *short* judgement and 2) a relatively *larger* stop-cue-related RPE transient, because the underlying subjective value was relatively low just before the stop-cue.

Now consider a particular (objective) time interval near the criterion time, for which the animal makes a mix of “long” and “short” choices (*e.g*., 1.74 s; Figure 4B). Soares et al. (2016) found that the amplitude of the stop-timing cue-related GCaMP6f transient tended to be bigger when the animal incorrectly made short choices, and this was taken as evidence that elevated DAN activity *slows* the internal clock. However, our model predicts that the size of the stop-cue-related transient will be inversely related to the amplitude of the underlying subjective value at that point, and thus inversely related to elapsed *subjective* time. It thus follows that if subjective time is more advanced on a given trial (*i.e*., faster pacemaker), the animal would tend to choose the long judgment on that trial, and the stop-timing RPE transient would be *smaller*. Conversely, if subjective time is less advanced on a trial (*i.e*., slower pacemaker), the animal would tend to choose the short judgment, and the stop-timing RPE transient would be *larger*.

Our RPE model accurately predicts the post-stop-cue dynamics of Soares et al. (2016); however, our model holds that elevated DAN activity *speeds* the internal clock, consistent with most pharmacological studies but *opposite* the interpretation of Soares et al. (2016). Thus, our RPE-based model suggests a parsimonious explanation for DAN activity in both the self-timed movement and temporal bisection paradigms, with (1) relatively high DAN activity corresponding to a fast pacemaker; manifesting in (2) compression of the value landscape; thereby leading to (3) early movements (in the self-timed movement task) or long judgments (in the temporal bisection task).

## Discussion

### Limitations of the RPE-based model

The simple RPE-based models presented here explain dynamic DAN signals in both the bisection task and our self-timed movement task, but they do not explain the origin of baseline offsets. Mohebi et al. (2019) recently-proposed that baseline offsets in ventral striatal dopamine levels could reflect the average recent reward rate, but we found that offset amplitude in DAN signals is at least partially independent of recent trial history during the self-timed movement task (Hamilos et al., 2021). It is possible that baseline variation arises from slow, random fluctuations in DAN activity, but further work is needed to explore the origins of these signals.

A second issue is the impact of optogenetic DAN activation and suppression on the rate of the pacemaker. In our self-timed movement task, DAN activation promoted early movements, consistent with increasing the pacemaker rate, whereas suppression promoted late movements, consistent with slowing the pacemaker rate (Figure 7 of Hamilos et al. (2021)). However, Soares et al. (2016) reported an opposite effect for optogenetic manipulation during the bisection task, at least for DAN activation.

This difference between the tasks could be reconciled by a recent theoretical model proposed by Mikhael and Gershman to explain the behavior of the pacemaker in a wide range of classical conditioning and timing studies (Mikhael and Gershman, 2019). Their model shows that the pacemaker rate is expected to be updated at the time of reinforcement by a Hebbian-like, bidirectional learning rule. If reward occurs exactly at the expected time, there is no update in the pacemaker rate. However, if reinforcement occurs before the expected time, this is interpreted as feedback that the pacemaker was running too slowly; thus, the update rule increases the pacemaker rate leading to expectation of reward at an earlier time on the next trial. Conversely, if reinforcement occurs after it was expected, this is interpreted as feedback indicating an overly fast pacemaker, resulting in an update that slows the pacemaker rate and creates the expectation of a later reward on the next trial. The same principles apply to ongoing RPE during timing tasks, and we recently showed that this model could predict trial-by-trial updates to the slope of ramping in our self-timed movement task (Jakob et al., 2022).

In our self-timed movement task, we activated or inhibited DANs only *up to* the time of first-lick, which Mikhael and Gershman’s model predicts will produce an effect on the pacemaker rate consistent with the sign of the manipulation (activate: increase, inhibit: decrease). However, Soares et al. (2016) continued optical stimulation *past* the end of the timing interval, until the end of the trial. When Mikhael and Gershman modeled stimulation in the Soares et al. (2016) task, they found that simulated DAN activation increased the pacemaker rate during the timing interval, but the continuing stimulation after the stop-timing cue rapidly counteracted this effect, resulting in *slowing* of the modeled pacemaker between the stop-cue and the judgment, leading to an effect on pacemaker rate *inconsistent* with the sign of the manipulation, as observed in Soares et al. (2016). If this model is correct, the effect of stimulation on the animal’s judgment in the Soares et al. (2016) task may have arisen due to continued manipulation of DAN activity *after* the timing interval had ended. A “retrospective” effect of this sort might seem counter-intuitive, but such retrospective effects have long been observed in perceptual studies, in which recall of sensory stimuli can be enhanced by additional sensory cues presented shortly after stimulus offset (Gegenfurtner and Sperling, 1993; Herrington and Assad, 2009), suggesting that sensory events are “buffered” briefly and can be altered by neural activity occurring between the sensory event and the perceptual decision. It is possible that a similar process could occur in the bisection task if DAN stimulation extends past the timing interval, although this is speculative. More work is needed to reconcile the optogenetic results in the self-timed movement and bisection tasks. To start, it would be informative to repeat the optogenetic experiments in the bisection task with optical stimulation limited to the period of the timed intervals only.

### Dopamine ramping as a population vs a single-cell signal?

Ramping signals in our photometry experiments were measured from a population of DANs. An important future question is whether ramps are also present at the level of individual neurons, or rather represent a progressive recruitment of individual neurons, or some combination of both. Prior studies have reported ramping signals in individual neurons during tasks with visual feedback of distance to reward (Kim et al., 2019), whereas others have observed decoupling between DAN firing rates and downstream dopamine release (Mohebi et al., 2019), making it unclear whether electrophysiology would be capable of addressing this question. Observation of individual neurons expressing calcium indicators with GRIN-lens equipped endoscopes may be better suited to addressing this question.

### Shape of the value function in behavioral timing

An intriguing feature to note is that DANs exhibited stretched, ramping dynamics both at cell bodies in SNc (Figures 1B and 3E, left) and VTA (Figure 3E, right), as well as in DAN axon terminals, and in striatal dopamine release (Hamilos et al., 2021). However, the stretched ramps differed qualitatively. In general, the medial dopaminergic ramping signals were the most convex and similar to the exponential form of the simulated learned value function in our model (Figure 3E, right).

Conversely, dopaminergic signals recorded more laterally in SNc had two distinct features: 1) ramps were more linear in quality, and, 2) ramps consistently showed an abrupt downturn starting about 300 ms before spout detection. This was earlier than the appearance of movement-related optical artifacts, which tended to occur within 150ms of spout contact. The middle panel of 3E shows averages for the 2 mice in our cohort who had the most lateral implants (which occured by chance due to variability in surgical placement and individual differences in anatomy). Intriguingly, DAN axon terminals recorded from dorsolateral striatum ubiquitously showed these down-turns as well in every mouse we recorded (n=7). These downturns are unlikely to be due to movement-related optical artifacts because a) they occur too early to align with the earliest inflections of our movement control channels, b) their amplitude is much larger than typical optical artifacts, and c) the direction of optical artifact excursions was inconsistent, whereas downturns were highly consistent (in dorsolateral striatum, 46% of optical artifact excursions were upward-going and 23% were neither consistently upward- or downward-going, with only 30% of transients consistently downward going.).

Recordings from SNc cell bodies in areas falling between these medial and lateral extremes showed gradual transition of ramping shapes, showing some convexity, but not as pronounced as was observed in VTA (see the SNc mouse shown in Figure 3E-left). The significance of these different ramping shapes remains uncertain, but could possibly reflect a process in which SNc or VTA neurons inherit information from one another to guide movement initiation. For example, VTA DANs may integrate SNc DAN ramps to signal reward expectation, an intriguing possibility given the recent report of dopamine waves in the literature (Hamid et al., 2021).

## Author Contributions

Allison Elizabeth Hamilos (A.E.H.) conceived the application and adaptation of the model to the two timing tasks, performed all experiments, implemented the simulations, and analyzed the data with John Abraham Assad (J.A.A). A.E.H. wrote the first draft of the manuscript, A.E.H. and J.A.A. finalized the paper.

## Methods

### Temporal difference learning under self-resolved state uncertainty

We adapted the temporal difference learning framework under state uncertainty resolved with visual feedback, which is described elegantly by Mikhael et al. (2022). There are some notable differences in our implementation for behavioral timing.

In Mikhael et al. (2022), state uncertainty is resolved by continual sensory feedback, as might be provided by visual cues during navigation. However, in behavioral timing, we assume that because the agent must rely on its subjective estimate of veridical time to guide behavior, it ”self-resolves” its state uncertainty continually with ”internally-generated feedback,” which is its subjective estimate of elapsed time. In other words, the internal timing signal is assumed to serve as evidence in the absence of external cues. Practically speaking, the implementation of this subtle difference is nearly the same as the original implementation by Mikhael et al. (2022). In both cases, it results in a continual reduction of state uncertainty at each state transition. Other differences in implementation are detailed below.

### Simulation details

#### Simulating traversal of subjective time states at different subjective pacemaker rates (Figures 2 and 3)

The self-timed interval was discretized into subjective timing states, *τ*_1_ : *τ_n_*, where each subjective timing state represented 80ms of veridical time spanning 4.4s. Thus there are n=43 time-states between the cue (defined as time *τ* = 0) and the first-lick state, *T*, which represents the optimal first-lick time-state at 3.36s. We assumed reward is not detected until the next state, *T* + 1, at 3.44s. We represented the lamp-off interval between the lamp-off and cue events as 5 time-states (0.4s total) before the cue, however we did not implement value function learning during this window. This could be applied in future models.

The simulation of subjective timing state traversal ran for each millisecond of veridical time. Within a subjective state, animals had a probability of exiting to the next timing state that depended on the speed of their internal clock. To model the behavior of the ideal timer in Figure 2, the probability of transitioning to the next subjective timing state was 1 when veridical time reached a multiple of 80ms (and 0 otherwise), such that the probability of transitioning at the end of any timing state was 1. To model variable speeds of the internal pacemaker, *p_txn_*was a number between (0, 1). In Figure 3, *p_txn_*(*slow*) = 0.008*/ms* and *p_txn_*(*fast*) = 0.0275*/ms* at each millisecond of the simulation. Because transitions were probabilistic, progression through the subjective timing states was simulated 1000 times to generate a 95% confidence interval for the average subjective timing state for each pacemaker rate.

The subjective timing state and the learned value and RPE functions (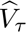 and *δ_τ_*, respectively) were subsequently mapped onto veridical time to simulate stretching/compression with differing pacemaker rates.

In this model, we assume the mouse has no information about its time-state before the cue event (though in the task, the animal *does* receive some information at the lamp-on event, which we have not accounted for here for simplicity. Future work with a more complex model employing lamp-off-to-cue interval time-states could potentially account for pre-cue dynamics.) We likewise did not aim to capture post-first-lick dynamics in this model, but rather focused on the timing interval. As such, models were plotted up to *T* for visualization.

#### Value landscape learning of an optimal behavioral timing agent (Figure 2)

We adapted the TD learning model for State Uncertainty Plus Feedback described by Mikhael et al. (2022) to the behavioral events of the self-timed movement task employed by Hamilos et al. (2021). We kept the learning parameters from the original model the same for consistency: *γ* = 0.9, *α* = 0.1, and Weber fraction *w* = 0.15, with animals iterating through each state on every trial and the learning iterated for 1000 trials. The standard deviation of the large uncertainty kernel before internal feedback was *σ_l_* = 3. After internal feedback, the standard deviation of the small uncertainty kernel was *σ_s_* = 0.1.

To avoid edge effects that had complicated the original model by Mikhael et al. (2022) (issues due to the value function being undefined after the first-lick), we assumed that uncertainty kernels existed out to effectively infinite time by modeling *n* = 63 total subjective timing states, such that max veridical time *t_n_*= 4.4*s*, which ensured kernels were fully-formed and well behaved at all simulated timepoints. Likewise, reward in the TD models was assumed to be 0 at all times except *r*(*T* + 1 : *T* + 10) = 1, which ensured learning of *V_h_* and *δ_τ_*was well behaved and not contaminated with artifacts from missing data to the right of the first-lick. The learned value function converged with good concordance to the ground truth value function using these parameters (Figure 2B).

#### Simulated value and RPE landscape of a perceptual timing agent (bisection task; Figure 5)

The value landscape was derived assuming a convex function during the timing interval with two abrupt increases, first at the tone 1 onset and once again at tone 2. Only the value landscape for the longest possible duration trial is shown for simplicity. For other duration trials, the value function was assumed to abruptly jump to its post-tone 2 trajectory following tone 2 onset. Following the principles of self-resolved state uncertainty, the RPE landscape for each trial duration was approximated as the derivative of the assumed value landscape by taking the difference 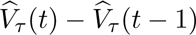 at every moment in time.

#### Simulating calcium indicator kinetics

Value and RPE curves were smoothed with a first-order Butterworth filter with frequency cutoff of 0.2Hz to simulate GCaMP6f kinetics.

### Visualization of calcium dynamics

#### Smoothing and trial pooling

Photometry signals were corrected for bleaching with dF/F, as detailed in the methods of (Hamilos et al., 2021). Calcium signals were pooled by movement time and plotted up to 150ms before movement in all figures. This was to reduce the possibility of contamination by optical artifacts, which typically occurred within 150ms of spout contact on average. For clarity of visualization, traces were smoothed with a 70ms gaussian smoothing kernel. Traces in Figure 1 were plotted as in the original paper, with trials pooled in equal 250 ms bins. In Figure 3, traces were pooled as follows: blue 1.5-3.3s; black 3.3-5.1s; red 5.1-7s (end of trial). Note that due to the shape of the movement distribution (Figure 1C inset), there are many more trials in the blue and black pools, and thus traces for longer movement times are expectedly noisier (i.e., the red traces). This arises from a different process than the wider 95% confidence band for red traces in Figure 3A-D, which occurs due to accumulated variability in the simulated state transition times at longer movement times as a result of the underlying probabilistic transition process.

## Notes

**Author note.** This work was supported by NIH grants UF-NS109177 and U19-NS113201, and NIH core grant EY-12196. A.E.H. was supported by a Harvard Lefler Predoctoral Fellowship, a Harvard Quan Predoctoral Fellowship, and a Harvard-MIT MD-PhD Program Scholarship. The content is solely the responsibility of the authors and does not necessarily represent the official views of the National Institutes of Health. The funders had no role in study design, data collection and analysis, decision to publish, or preparation of the manuscript.

https://github.com/harvardschoolofmouse/RPE-Modeling

### Competing Interest Statement

The authors have declared no competing interest.

### Summary of Updates

We have expanded upon behavioral timing models from the first paper, including new data. We have crested a GitHub repository with the behavioral timing models. We have added a methods section and extended discussion.

